# *Plasmodium falciparum* GCN5 plays a key role in regulating artemisinin resistance–related stress responses

**DOI:** 10.1101/2023.01.11.523703

**Authors:** Amuza Byaruhanga Lucky, Chengqi Wang, Ahmad Rushdi Shakri, Mohammad Kalamuddin, Anongruk Chim-Ong, Xiaolian Li, Jun Miao

## Abstract

*Plasmodium falciparum* causes the most severe malaria and is exposed to various environmental and physiological stresses in the human host. Given that GCN5 plays a critical role in regulating stress responses in model organisms, we aimed to elucidate PfGCN5’s function in stress responses in *P. falciparum*. The protein level of PfGCN5 was substantially induced under three stress conditions (heat shock, low glucose starvation, and dihydroartemisinin, the active metabolite of artemisinin (ART)). With a TetR-DOZI conditional knockdown (KD) system, we successfully down-regulated PfGCN5 to ∼50% and found that KD parasites became more sensitive to all three stress conditions. Transcriptomic analysis via RNA-seq identified ∼1,000 up-and down-regulated genes in the wildtype (WT) and KD parasites under these stress conditions. Importantly, DHA induced transcriptional alteration of many genes involved in many aspects of stress responses, which were heavily shared among the altered genes under heat shock and low glucose conditions, including ART-resistance-related genes such as *K13* and *coronin*. Based on the expression pattern between WT and KD parasites under three stress conditions, ∼300-400 genes were identified to be involved in PfGCN5-dependent, general and stress-condition-specific responses with high levels of overlaps among three stress conditions. Notably, using ring-stage survival assay (RSA), we found that KD or inhibition of PfGCN5 could sensitize the ART-resistant parasites to the DHA treatment. All these indicate that PfGCN5 is pivotal in regulating general and ART-resistance-related stress responses in malaria parasites, implicating PfGCN5 as a potential target for malaria intervention.

**IMPORTANCE:** Malaria leads to about half a million deaths annually and these casualties were majorly caused by the infection of *Plasmodium falciparum*. This parasite strives to survive by defending against a variety of stress conditions, such as malaria cyclical fever (heat shock), starvation due to low blood sugar (glucose) levels (hypoglycemia), and drug treatment. Previous studies have revealed that *P. falciparum* has developed unique stress responses to different stresses including ART treatment, and ART-resistant parasites harbor elevated stress responses. In this study, we provide critical evidence on the role of PfGCN5, a histone modifier, and a chromatin coactivator, in regulating general and stress-specific responses in malaria parasites, indicating that PfGCN5 can be used as a potential target for anti-malaria intervention.

## INTRODUCTION

Malaria is one of the most severe public health problems worldwide. *Plasmodium falciparum* causes the most severe form of malaria and is responsible for about half a million deaths annually [1]. Malaria parasites are exposed to various environmental and physiological stresses in the human host (e.g., cyclical fever, low nutrition, and drug treatment) [2]. They have evolved general and specific mechanisms to defend against those assaults [3–5]. Several chaperones (PfHSP70-1, PfHSP70-x, PfHSP110, and the endoplasmic reticulum chaperone PfGRP170) were identified as essential proteins for the parasite to respond to heat shock (HS) [6–10]. PfAP2-HS, an ApiAP2 (AP2) domain-containing transcription factor, was found to rapidly activate *Pfhsp70-1* and *Pfhsp90* in the protective HS response [5]. Besides the conserved mechanisms for tolerance to febrile temperature and oxidative stress such as redox and protein-damage responses, parasites developed specific mechanisms such as regulating isoprenoid biosynthesis and its downstream protein modifications (geranylgeranylation and farnesylation) [4, 11]. Isoprenoid biosynthesis occurs in the apicoplast, an apicomplexan pathogen-specific organelle derived from an algal endosymbiont plastid. Many genes targeting the apicoplast were also upregulated upon HS, suggesting that the parasite utilizes an analogous defense system against heat stresses like plants [4, 12, 13].

Malaria parasites employ similar mechanisms to deal with the stresses of antimalarial treatment. ART, the first line of the anti-malaria drug, causes the production of free radical species, including reactive oxygen species (ROS), which damage proteins, lipids, and DNA [14–25]. Low-dose ART treatment was found to activate the response pathways critical for the tolerance to febrile temperature [4]. ART-resistant parasites with mutations in the Kelch protein 13 (PfK13) up-regulated genes related to stress responses (e.g., protein folding, redox, and proteasome-linked protein turnover). ART-resistant parasites respond to ART treatment by elevating gene expression related to apicoplast and mitochondrial metabolism, vesicular trafficking, lipid transport, and tRNA modifications [21–25].

GCN5 is a well-known key regulator of stress responses in humans, plants, yeast, and *Toxoplasma* by coordinating with specific transcriptional factors [26–33]. Recent studies also identified up-regulation of PfGCN5 in response to stress conditions (ART treatment, glucose starvation, and HS) along with many other up-regulated genes in *P. falciparum* [3, 34]. Intriguingly, PfGCN5 was found to bind many of these genes, but most of the binding sites were localized in the coding regions, not in promoters [35]. Treatment with garcinol, a PfGCN5 inhibitor, sensitized the ART-resistant *P. falciparum* parasite to ART during the ring stage. In addition, treating parasites with ART caused substantial changes in the abundance of active chromatin markers H3K9ac and H4K8ac [35]. Collectively, these studies provided tangential evidence implying PfGCN5’s participation in responses to ART.

To elucidate the functions of PfGCN5 in orchestrating the transcriptional program in *P. falciparum*, we deleted the C-terminal bromodomain of PfGCN5, which supposedly mediates the binding of PfGCN5 to acetylated lysines in histone. This led to drastic transcriptional changes in many genes, including protein folding-related genes and *AP2-HS* [36], suggesting that PfGCN5 regulates the stress response pathways. However, this assumption is undermined by the dislocation of the PfGCN5 complex from its chromatin targets due to bromodomain deletion. To clarify the critical role of PfGCN5 in regulating stress responses in *P. falciparum*, we employed a conditional knockdown (KD) system to down-regulate PfGCN5 and determined the parasite’s responses to different stress conditions. We provide critical evidence about the role of PfGCN5 in regulating general and stress-specific responses in malaria parasites, including the stress responses in ART resistance, implicating PfGCN5 as a potential target for therapeutic development.

## RESULTS

### Conditional KD of PfGCN5 impairs parasite growth

PfGCN5 is essential for asexual blood stages, and deletion of the C-terminal bromodomain led to a defect in RBC invasion and dysregulated expression of virulence genes [36]. To elucidate whether and how PfGCN5 regulates stress responses, we employed the TetR-DOZI system [37, 38] to conditionally knock down PfGCN5 expression. We created a parasite line, TetR-PfGCN5::GFP, with the insertion of the 10× aptamer in the 3’ end of the endogenous PfGCN5 locus and fusion of the GFP tag to the PfGCN5 C-terminus, which would allow us to monitor PfGCN5 expression (**Figure 1A, S1A**). Correct integration of the plasmid at the *PfGCN5* locus was confirmed by a genomic Southern blot (**Figure S1B**). The binding of the TetR-DOZI to the aptamer in the presence of anhydrous tetracycline (+aTc) allowed the expression of PfGCN5-GFP. Western blot showed that PfGCN5-GFP in the TetR-PfGCN5::GFP line was expressed at a similar level and was proteolytically processed into the same five fragments compared to PfGCN5-GFP in the published PfGCN5::GFP line and native PfGCN5 by anti-PfGCN5 antibodies [36, 39] (**Figure S1C**). Withdrawal of aTc (−aTc) for one intraerythrocytic developmental cycle (IDC) led to ∼50% reduction of PfGCN5-GFP expression, shown in GFP fluorescence intensity by flow cytometry analysis (**Figure 1A, 1B**). Intriguingly, the withdrawal of aTc beyond one IDC did not further reduce the PfGCN5 expression level (**Figure S1D**), suggesting that parasites were probably restrained from further reduction of PfGCN5 expression due to its essentiality. Western blots showed the same level of reduction (∼50%) in full-length PfGCN5 protein as well as its cleaved fragments after KD (**Figure 1C**). Live cell imaging revealed that PfGCN5-GFP in the TetR-PfGCN5::GFP was localized in the parasite nucleus like the PfGCN5::GFP parasite in our earlier study [36] and KD resulted in dimer GFP signals in the nucleus (**Figure 1D**). The growth rate of −aTc parasites was notably slower than that of the +aTc parasites starting from the second cycle (**Figure 1E**). We found that PfGCN5 KD was reversible, as within ∼8-10 h of aTc add-back, PfGCN5-GFP expression returned to the level of the +aTc culture determined by flow cytometry and Western blots (**Figure S1D, S1E**), and the growth rate of the parasites was rapidly restored (**Figure S1F**).

**Figure 1.**
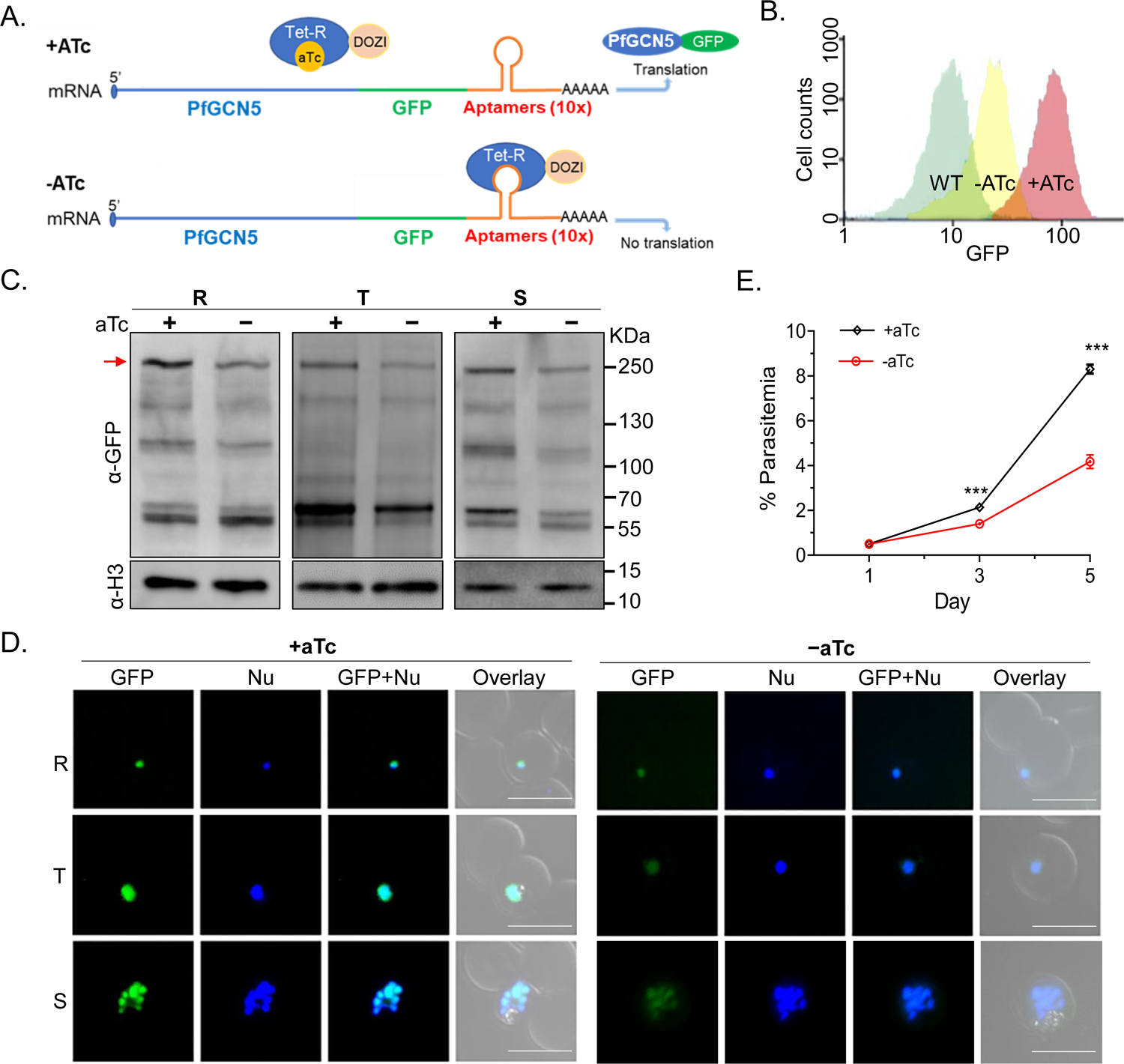
KD of PfGCN5 led to parasite growth defects and increased susceptibilities to stress conditions. **A**. A diagram illustrates the TetR-DOZI inducible KD system, where 10x aptamer motifs are inserted into the 3’ UTR of the *PfGCN5*. Adding aTc causes TetR-DOZI to be released from the aptamer, inducing protein translation (translation “ON”), whereas withdrawal of aTc leads to the binding of TetR-DOZI to the aptamer motifs to block the translational process (translation “OFF”), “AAAAA” indicates the polyA tail of the mRNA. **B**. Measurement of GFP expression in the TetR-PfGCN5:GFP parasites by flow cytometry. The reduction of PfGCN5-GFP protein level after withdrawal of aTc for 48 h is shown. **C**. Western blot with anti-GFP antibodies (α-GFP) showed the reduction of PfGCN5-GFP after PfGCN5 KD by the withdrawal of aTc at the ring (R), trophozoite (T), and schizont (S) stages for five IDCs. The histone H3 was used as a loading control. The red arrow indicates the positions of the full-length bands of PfGCN5-GFP. **D**. The growth curves of TetR-PfGCN5:GFP parasites with or without aTc. The parasite growth rates were significantly reduced after KD of PfGCN5 (−aTc) compared to the parasites without KD of PfGCN5 (+aTc) (*p* < 0.001, multiple T-test). **D**. GFP fluorescence signals of PfGCN5-GFP in the TetR-PfGCN5:GFP parasites at the ring (R), trophozoite (T), and schizont (S) stage before (+aTc) and after PfGCN5 KD by the withdrawal of aTc (−aTc) for five IDCs. Nu: nuclear staining of live parasite-infected RBCs by Hoechst 33342. The size of the scale bar is 10 μm.

### Stress induces PfGCN5 expression and PfGCN5 KD leads parasites more sensitive to stress

Previous studies showed that 6h of stress conditions (HS, low glucose, and ART treatment) could substantially induce the transcriptional expression of PfGCN5 in the early-stage parasites (22 hours post-invasion, hpi) by quantitative reverse transcription PCR (RT-qPCR) [3, 34]. To determine whether PfGCN5 could be induced at the protein level, we performed Western blots to measure the PfGCN5 proteins in the stressed parasites. After the TetR-PfGCN5::GFP parasite line cultured with aTc was treated at the early ring stage (0-6 hpi) with three stress conditions: HS (41 °C for 6 h), low glucose (0.5 g/L for 6 h), or DHA (30nM for 6 h), PfGCN5-GFP protein was significantly induced with a surprising finding that starvation caused the most significant induction of PfGCN5 expression (**Figure 2A**). A similar trend was found after the same stress treatments to the late stage (trophozoite, 24-30 hpi) of the TetR-PfGCN5::GFP parasite line (**Figure S2A**).

**Figure 2.**
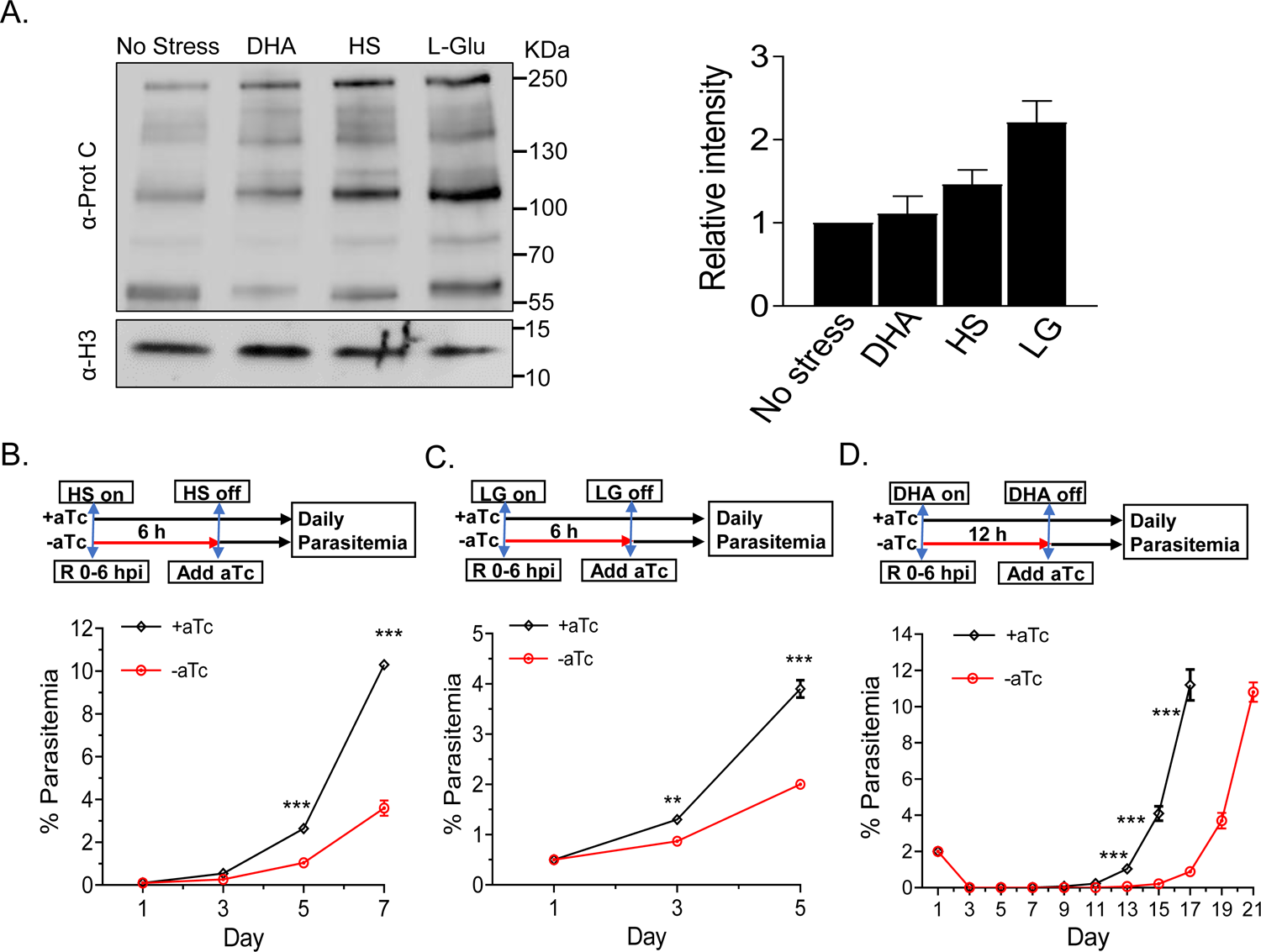
The expression of PfGCN5 and parasite growth upon stress treatments. **A**. The left panel shows the changes of PfGCN5-GFP expression after +aTc TetR-PfGCN5:GFP parasites were treated with heat shock (HS, 41 °C), low-glucose (LG, 0.5 g/L), and DHA (30mM) for 6 h at early ring stage by Western blot. The histone H3 was used as a loading control. The right panel indicates the relative intensity of full-length PfGCN5-GFP bands among the parasites with or without stress treatments by densitometry. **B** and **C**. Parasite growth with or without *PfGCN5* KD exposed to HS (**B**) and low glucose (LG) (**C**) conditions, respectively. (**: *P* < 0.01, ***: *P* < 0.001, multiple T-test). TetR-PfGCN5:GFP parasites at the early ring stage with (+aTc) or without aTc (-aTc) were treated by HS (41 °C) and low-glucose (0.5 g/L) for 6 h. aTc was added back to the culture after stress treatment. **D**. Parasite recovery assays showing the time required for the control (+aTC) or the PfGCN5 KD (-aTc) parasites to recover after treatment with 1 μM of DHA for 12 h (*P* < 0.001, multiple T-test). aTc was added back to the culture immediately after DHA treatment.

To investigate if PfGCN5 is involved in regulating stress responses, the TetR-PfGCN5::GFP parasite line cultured with or without aTc was treated at the early ring stage (0-6 hpi) with different stress conditions: HS (41 °C for 6 h), low glucose (0.5 g/L for 6 h), or DHA (1 μM for 12 h) (**Figure 2B-D**). After removing the stress conditions, aTc was added back to the culture to restore the expression of PfGCN5 and exclude the impact of PfGCN5 KD on the parasite growth after stress treatment and parasite growth was monitored daily. TetR-PfGCN5::GFP parasites without aTc grew significantly more slowly than +aTc parasites under HS and low-glucose conditions (**Figure 2B, 2C**). After DHA treatment, both +aTc and −aTc parasites were non-detectable through day 9. The +aTc culture resumed growth and reached 5% parasitemia on day 15, whereas the −aTc culture had a 4-day delay in reaching 5% parasitemia (**Figure 2D**). This delayed growth phenotype was not caused by aTc because the 3D7 WT parasite treated the same way showed the same growth pattern after DHA treatment (**Figure S2B**). Taken together, PfGCN5 KD reduced the parasite’s tolerance to different stresses, strongly suggesting a direct association between PfGCN5 and the regulation of stress responses.

### Parasites apply general and specific responses to different stress conditions

To determine how the parasites respond to different stress conditions, we first characterize the transcriptomic changes of TetR-PfGCN5::GFP parasites (+aTc) at the ring-stage (0-6 hpi) after treatment with HS (41°C), low glucose (0.5 g/L), or DHA (30 nM) for 6 h. Transcriptomic analysis was performed by RNA-seq with three biological replicates. Pearson correlation and principal component analysis (PCA) indicated that there was high consistency among the replicates (**Table S1, Figure S3A**). DESeq2 analysis using a cutoff of *P-adj* < 0.1 and > 1.5-fold [40] identified 1183, 1130, and 1038 up-regulated, as well as 1151, 1126, and 1049 down-regulated genes by HS, low-glucose, and ART treatment, respectively (**Figure 3A-C, Table S1**). Surprisingly, there were substantial overlaps in the up-and down-regulated genes among different stress conditions (57-65% in the up-regulated genes and 70-76% in the down-regulated genes) (**Figure 3D and E, Table S1**). Gene ontology (GO) enrichment analysis based on biological process (BP) (**Figure 3F**) and cellular component (CC) (**Figure S3B**) showed that genes related to translation and tRNA metabolism, protein-damage responses (protein folding and proteasome), glycolysis and gluconeogenesis, nucleotide metabolism, host-cell remodeling, and mitochondrial and apicoplast proteins were up-regulated in all stress conditions. In contrast, genes related to merozoite invasion, egression, DNA replication, cytoskeleton, protein phosphorylation, and phospholipid transport were downregulated, indicating that parasites used a general stress response to different stress conditions (**Figure 3G, S3C**).

**Figure 3.**
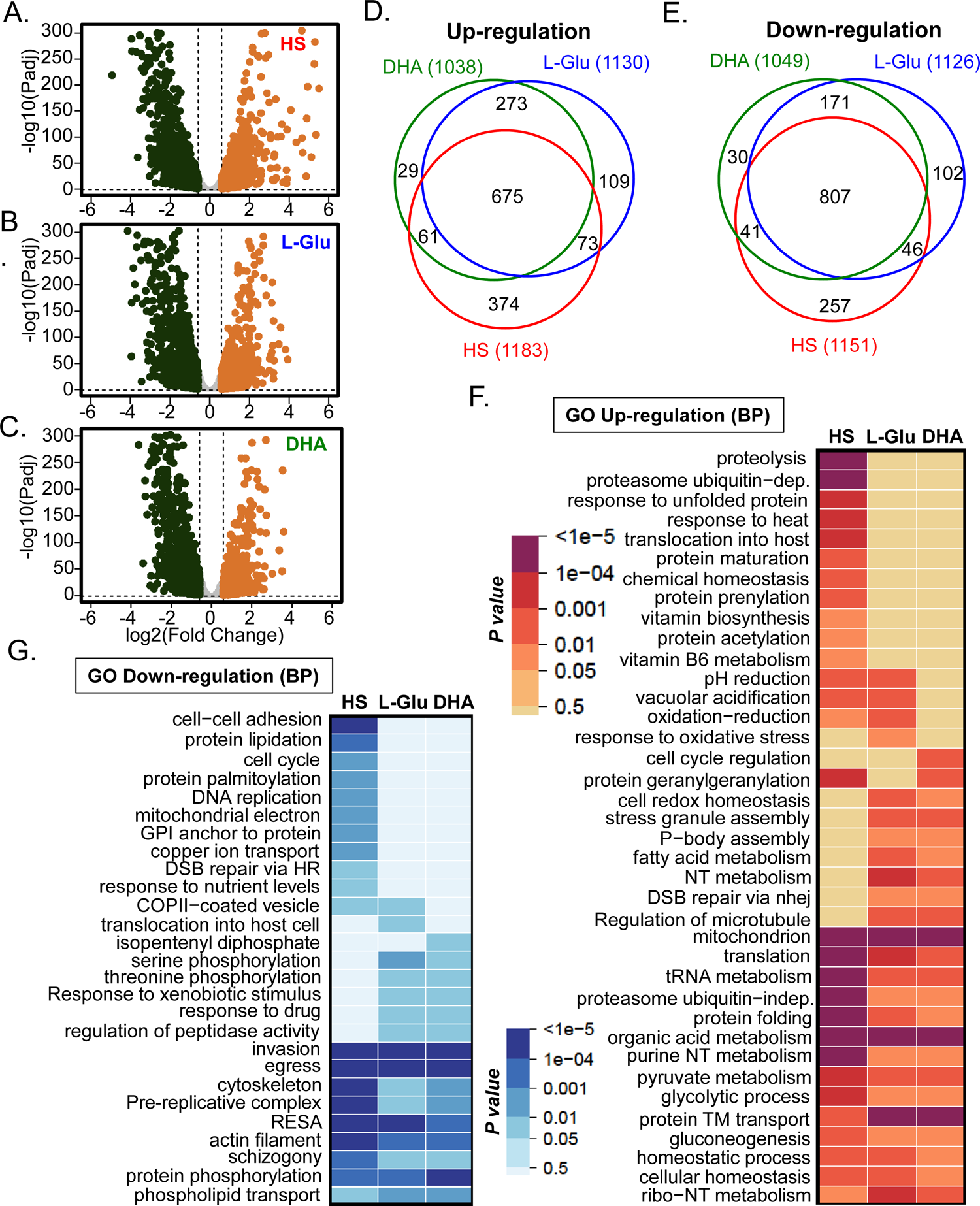
Transcriptional changes of +aTc TetR-PfGCN5::GFP parasites upon exposure to different stress conditions. **A**-**C**. Volcano plots showing differentially expressed genes at the ring stage after treatment with heat shock (HS) (**A**), low glucose (L-Glu) (**B**), and DHA (**C**). **D** and **E**. Venn diagrams show the number of altered genes and the extent of overlaps among the up-(**D**) and down-(**E**) regulated genes upon HS, low glucose, and DHA treatments, respectively. **F** and **G**. Heatmaps display the GO enrichment analyses of up-(**F**) and down-(**G**) regulated genes upon HS, low glucose, and DHA treatments based on the biological process (BP), showing the common and stress-condition-specific stress responses.

The transcriptomic analysis also revealed many genes whose expression was altered in a stress condition-specific manner (**Figure 3F, 3G, S3B, and S3C**). HS specifically induced the up-regulation of genes related to the response to heat and protein unfolding, including *Pfhsp70-1* and *Pfhsp90*, which are regulated by the transcription factor PfAP2-HS [5]. The low-glucose condition specifically activated genes related to oxidative stress response and mitochondrial ATP synthesis. ART treatment specifically up-regulated genes related to cell-cycle regulation and ER stress response. Genes related to protein geranylgeranylation (apicoplast function) were only enriched after HS and ART treatment but not under glucose starvation (**Figure 3F**, **Table S1**).

ART resistance in *P. falciparum* was found to be medicated by mutations in the propeller domain of PfK13 [41], and the expression levels of PfK13-interacting proteins (KICs) were found to influence hemoglobin uptake [42]. Intriguingly, PfK13, three kelch domain-containing proteins (PF3D7_1125800, PF3D7_1125700, and PF3D7_0724800), seven of ten KICs (KIC1-3, 5-8) were significantly down-regulated in all three stress conditions, whereas KIC4 was significantly down-regulated under low glucose and KIC10 was down-regulated under HS and low glucose conditions. Although the *P-adj* values for KIC10 (0.000194) downregulation under DHA treatment and KIC4 downregulation under DHA (1.09e-10) and HS (0.00156) was substantially lower than the cutoff of *P-adj* (<0.1), the fold change for KIC10 (1.447), and KIC4 (DHA:1.3005 and HS:1.1532) did not reach the cutoff of 1.5, respectively (**Table S1**). Another ART resistance-related gene, coronin (PF3D7_1251200) [43, 44], was also significantly down-regulated in all three stress conditions. Additionally, four autophagy-related genes (ATG-5, -8, -11, and -23) were also significantly down-regulated in all three stress conditions. 27 AP2-TFs were transcriptionally altered with the same trends upon three stress conditions except that four AP2-TFs (PF3D7_0516800, PF3D7_0404100, PF3D7_1456000, and PF3D7_1239200) were up-regulated under DHA and low-glucose conditions but downregulated under HS (**Figure S3D**). AP2-HS, a regulator of HS response, was found upregulated at the highest level (1.3278) compared to other conditions (DHA: 1.152 and low-glucose: 1.0615) (**Table S1**). Collectively, these data indicate that different stress conditions induce general and stress-condition-specific stress responses in malaria parasites.

### PfGCN5 plays a crucial role in the regulation of stress responses

To understand how PfGCN5 is involved in stress responses, we sought to determine the transcriptomic changes of the parasites after manipulating PfGCN5 expression in response to different stress conditions. Transcriptomic analysis of the TetR-PfGCN5::GFP parasite at the early ring stage identified only two genes (*PfGCN5* and *EBA175*) that were significantly down-regulated (> 1.5-fold, *P-adj* < 0.1) after PfGCN5 KD (−aTc), indicating that reduced PfGCN5 expression did not disturb the overall transcription program at the early ring stage (**Table S2**). We then subjected the −aTc TetR-PfGCN5::GFP parasites (at least cultured without aTc for five IDCs) at 0-6 hpi to the same stress conditions for +aTc TetR-PfGCN5::GFP parasites mentioned above for 6 h and harvested RNA for RNA-seq analysis. Similar to the transcriptomes of +aTc TetR-PfGCN5::GFP parasites, Pearson correlation and PCA indicated that there was high consistency among the replicates of −aTc TetR-PfGCN5::GFP parasites (**Table S2, S3, Figure S3A**). In the −aTc TetR-PfGCN5::GFP parasites, the stress conditions HS, low glucose, and ART treatment resulted in a similar number of genes with expression changes as in the +aTc parasites: 971, 1136, and 911 up-regulated genes and 922, 1170, and 1041 down-regulated genes, respectively (**Figure 4A, 4B, Table S3**). These transcriptionally altered genes under three different stress conditions also overlapped substantially, with 48-60% and 66-84% overlaps among the up-and down-regulated genes, respectively (**Figure 4A, 4B**, **Table S3**).

**Figure 4.**
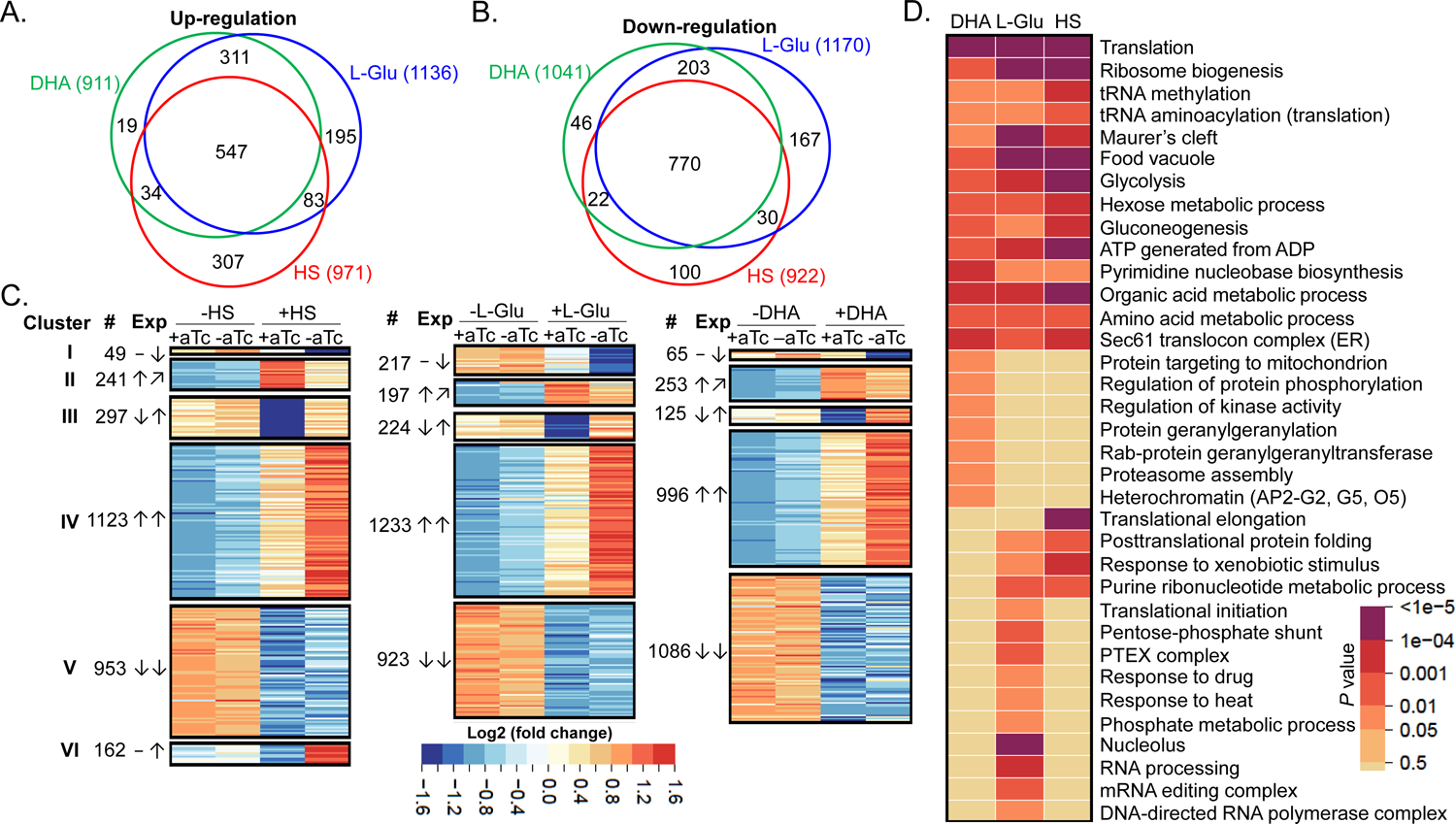
Transcriptomic analyses of PfGCN5-dependent stress responses. **A** and **B**. Venn diagrams showing the number of altered genes and the overlaps among the up- (**A**) and down- (**B**) regulated genes upon DHA, HS, and low-glucose treatments in the −aTc (PfGCN5 KD) parasites, respectively. **C**. Heatmaps show the expression patterns (Exp) in WT (+aTc) and PfGCN5 KD (−aTc) parasites upon HS, low glucose (L-Glu), and DHA treatments. ↑, ↓, and − denote genes that were up-, down-, and not-altered after treatments, ↗ indicates the genes failed to be up-regulated in PfGCN5 KD parasite after ART treatment. #: numbers of genes in each cluster. **D**. GO enrichment analyses of PfGCN5-dependent, stress response genes (cluster I and II) based on the *P* values.

Comparison of the transcriptomes between the −aTc and +aTc parasites allowed us to identify 2825, 2738, and 2528 genes that were differentially regulated by the HS, low glucose, and DHA treatment, respectively (**Table S4**). Based on their expression patterns, these genes were grouped into five clusters (**Figure 4C, Table S4**), while HS included an additional cluster (VI). Cluster I and II genes were down-regulated and failed to be up-regulated in −aTc parasites compared to +aTc parasites under three stress conditions, respectively, suggesting that these two clusters are likely the PfGCN5-dependent stress-response genes (**Figure 4C**). GO enrichment analysis showed that those genes in clusters I and II under three stress conditions are commonly involved in many aspects of critical pathways such as translation (ribosome biogenesis and tRNA modification), energy metabolism (ATP metabolism, glycolysis, hexose metabolic process, and gluconeogenesis), Pyrimidine and amino acid metabolic process, Sec61 translocon complex (ER-associated degradation (ERAD)), protein exported beyond parasite (Maurer’s cleft), and digestion (food vacuole) (**Figure 4D**). Furthermore, different stress also led to stress condition-specific, PfGCN5-dependent stress-response (**Figure 4D**). DHA treatment-specific, PfGCN5-dependent stress-response genes are involved in protein geranylgeranylation, regulation of protein phosphorylation/kinase, proteasome assembly, protein targeting mitochondrion and heterochromatin (AP2-G2, -G5, and -O5). The low glucose condition-specific, PfGCN5-dependent stress-response genes are specifically related to translation initiation, pentose-phosphate shunt (a major regulator for cellular reduction-oxidation (redox) homeostasis and biosynthesis), PTEX complex, response to drug and heat, and RNA process (nucleolus, mRNA editing, and RNA polymerase). Likewise, the heat shock-specific, PfGCN5-dependent stress-response genes are specifically involved in translational elongation. More genes related to protein folding and response to the xenobiotic stimulus were identified in the heat shock-specific, PfGCN5-dependent stress-response gene list than the ones in the low glucose condition-specific genes. Notably, KIC4, which was significantly down-regulated only after glucose starvation in +aTc parasites, was significantly down-regulated in −aTc parasites under all three stress conditions. Similarly, GARP (glutamic acid-rich protein) expressed on the surface of parasite-infected RBC [45, 46] also failed to be up-regulated in −aTc parasites under three stress conditions (**Table S4**). Antibodies against GARP killed the parasites and were positively associated with protection against severe malaria in children [47].

Conversely, cluster III genes, down-regulated in the +aTc parasites, were up-regulated in the −aTc parasites under three stress conditions, probably to compensate for PfGCN5 KD (**Figure S4A**). These include genes related to DNA replication and repair, cell cycle, and isoprenoid biosynthesis. Similarly, some genes were up-regulated only by glucose starvation (cell adhesion and cytoskeleton) and HS (mitochondrion electron transport). A large number of genes belong to clusters IV and V, which were up- and down-regulated in the −aTc parasites under stress conditions in the same trends as in the +aTc parasite, respectively, suggesting that these genes may be PfGCN5-independent. HS induced a new cluster of genes (VI), which were up-regulated in the −aTc parasites but no change in the +aTc parasites (**Figure 4C**). These genes are involved in similar pathways as the cluster III genes, suggesting that HS might have involved more genes in compensation for PfGCN5 KD (**Table S4**). By comparing the overlaps within these clusters upon three different stress conditions, low levels of overlaps were found among PfGCN5-dependent stress-response genes (cluster I and II) and the KD compensation-related genes (cluster III), whereas high levels of overlaps among the PfGCN5-independent stress-response genes (cluster IV and V) were identified (**Figure S4B, Table S4**), suggesting PfGCN5-dependent stress responses are also stress-condition-specific to a certain degree. Taken together, these data demonstrate that PfGCN5 regulates stress response by targeting stress response genes that are general and stress condition-specific to stress conditions.

To confirm that PfGCN5 regulates these PfGCN5-dependent, stress-response genes, we investigated the enrichment of PfGCN5 at the promoter regions (5’ UTRs) of three shared, PfGCN5-dependent genes under three stress conditions by chromatin-immunoprecipitation quantitative PCR (ChIP-qPCR). These three genes encode KIC4, carbamoyl phosphate synthetase (cpsSII) for de novo pyrimidine synthesis, and karyopherin beta (KASbeta) for nuclear import, respectively (**Figure 5**). We designed three pairs of primers for each gene to amplify the enrichment signals in the 5’ UTRs. As we expected, there were no significant differences in PfGCN5 enrichment between the WT (+aTc) and PfGCN5 KD (−aTc) when the parasites were not under stress conditions at the early ring stage whereas, under the stress conditions, PfGCN5 enrichment in WT (+aTc) was significantly higher than the one in PfGCN5 KD (−aTc), confirming that PfGCN5 KD led to the defect in activating these genes under stress conditions.

**Figure 5.**
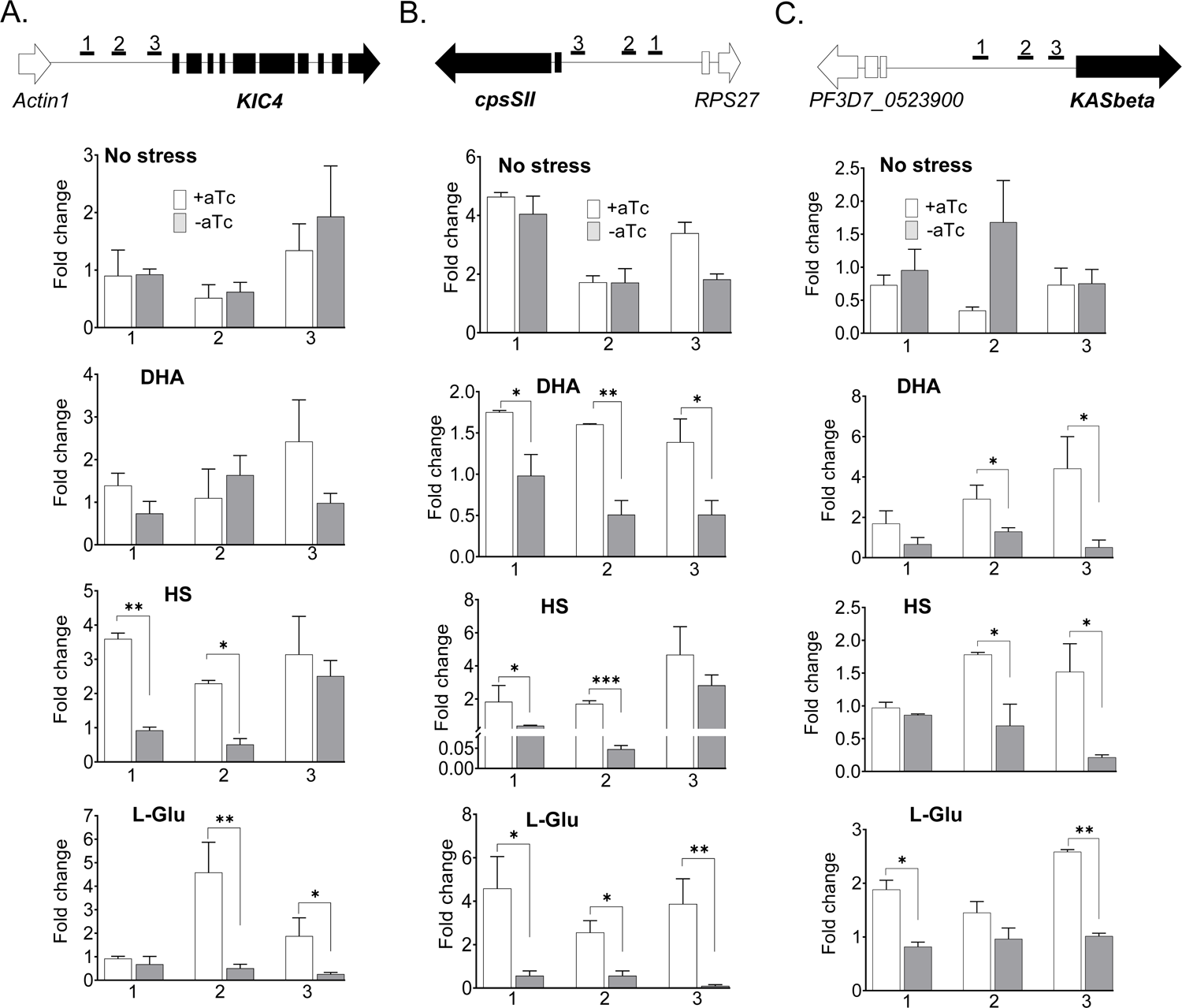
PfGCN5 directly regulates stress response genes by targeting 5’ UTRs. The enrichment of PfGCN5 was determined at the ring stage by ChIP-qPCR. **A-C.** The top panel shows the schematics of genomic loci and primer pairs marked as 1, 2, and 3 located in the 5’ UTR regions of *KIC4* (**A**), *cpsSII* (**B**), and *KASbeta* (**C**). The second to fifth panels show that ChIP-qPCR detected the enrichment of PfGCN5 in the 5’ UTR regions of the selected genes in the TetR-PfGCN5::GFP parasites with (+aTc) or without aTc (−aTc) under no stress (second), DHA (third), HS (fourth), and low glucose (L-Glu) condition for 6 h. The fold change indicates the enrichment relative to the reference gene *seryl-tRNA synthetase* (PF3D7_0717700). *, **, and *** indicate *P* < 0.05, 0.01, and 0.001, respectively, Mann-Whitney U test.

### KD or inhibition of PfGCN5 reduces parasites’ tolerance to DHA treatment

Growth recovery assay and transcriptomic analysis after DHA treatment revealed PfGCN5’s involvement in regulating general and specific responses to DHA (**Figure 2D, 4D**). To translate this phenotype into ART sensitivity, we performed the ring-stage survival assay (RSA) with TetR-PfGCN5::GFP parasites. The TetR-PfGCN5::GFP (−aTc) parasites at the early ring were exposed to 700 nM DHA for 6 h, and aTc was added back right after DHA treatment to exclude the subsequent effect of PfGCN5 KD on parasite growth. Compared to the RSA value of the TetR-PfGCN5::GFP parasites under the +aTc conditions, PfGCN5 KD resulted in a ∼60% reduction in the RSA value (**Figure 6A**).

**Figure 6.**
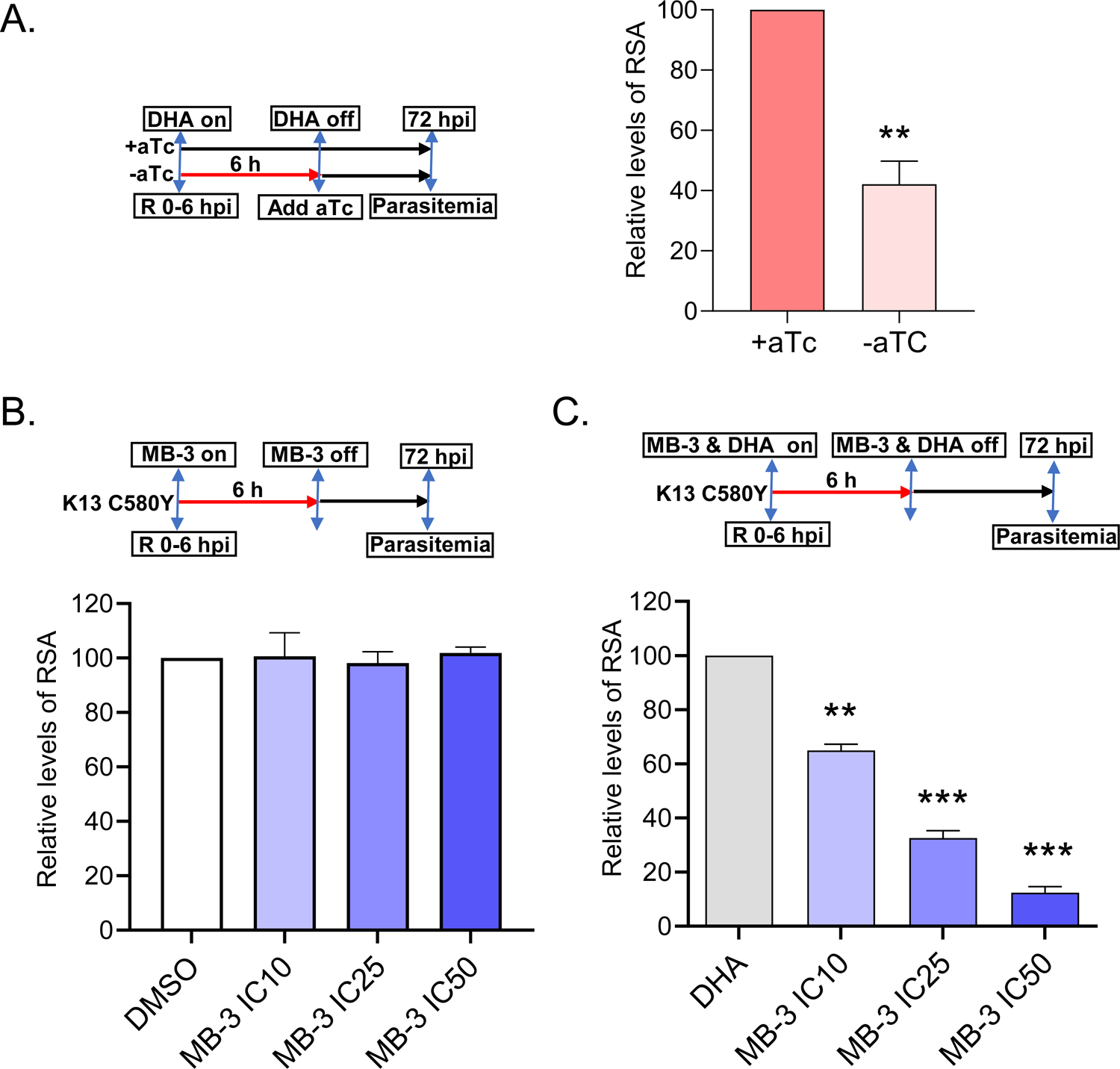
PfGCN5 KD or PfGCN5 inhibitor sensitized the parasites to DHA treatments. **A**. The left panel indicates the procedure of the RSA experiment. aTc was added back to the TetR-PfGCN5::GFP culture right after 6 h treatment by 700nM DHA. The right panel shows the RSA results of +aTc and -aTc parasites (TetR-PfGCN5::GFP). KD of PfGCN5 by the withdrawal of aTc (-aTc) led to the reduction of RSA values (*P* < 0.01, T-test). RSAs of +aTc parasite (∼0.3%) were set up as 100% for comparison. **B** and **C**. The top panels show the procedure of RSA experiments. RSA of an ART-resistant K13 mutant strain (isolated from Cambodia with C580Y mutation) while MB-3 was added and withdrawn at the same time as DHA treatment. The bar graphs indicate the results of RSA. MB-3 at the concentrations of IC_10_ (10.9 μM), IC_25_ (17.3 μM), and IC_50_ (27.5 μM) without DHA did not cause any noticeable alteration of RSA value as compared to DMSO vehicle control (**B**) whereas co-incubation with both DHA and MB-3 significantly reduced RSA values compared to DHA treatment only (**C**) (**: *P* < 0.01, ***: *P* < 0.001, T-test).

With the demonstration of PfGCN5’s role in regulating parasite’s responses to ART drugs, we wanted to test if inhibition of PfGCN5’s enzymatic activity would re-sensitize ART-resistant parasites to ART drugs. We evaluated butyrolactone 3 (MB-3), which we showed previously to inhibit PfGCN5’s enzymatic activity with an IC_50_ of ∼125 μM [48]. With the standard SYBR green I growth inhibition assay, MB-3 inhibited 3D7 parasites at an IC_50_ of ∼27.5 μM. We found that a 6-h exposure of Cam2, an ART-resistant strain collected from Cambodia with the K13 C580Y mutation (MRA-1236), at the early-ring stage to 10.9, 17.3, and 27.5 μM of MB-3, corresponding to the IC_10_, IC_25_, and IC_50_ concentrations of the drug, respectively, did not cause noticeable changes in the survival rate of ART-resistant strain as compared to the DMSO vehicle control (**Figure 6B**). Whereas RSA of Cam2 showed a survival rate of 13%, co-incubation of DHA and MB-3 for 6 h significantly reduced the RSA value in an MB-3 concentration-dependent manner compared to DHA treatment only (**Figure 6C**). Taken together, these data showed that KD or inhibition of PfGCN5 could re-sensitize the ART-resistant parasites to ART drugs.

## DISCUSSION

This study aimed to confirm PfGCN5’s involvement in regulating stress responses in the malaria parasite and gain a mechanistic understanding of PfGCN5’s role in responding to different stress conditions. Using a conditional KD system, we successfully down-regulated PfGCN5 expression and showed that PfGCN5 KD increased the parasite’s susceptibility to all stress conditions used, emphasizing PfGCN5’s central role in stress response. Through transcriptomic analysis, we identified 300-400 genes involved in PfGCN5-dependent, general, and stress-specific responses. Furthermore, using growth recovery assay and RSA, we found that KD or inhibition of PfGCN5 could sensitize the ART-resistant parasites to the ART treatment.

We have shown that the malaria parasites can mount a rapid stress response, with the expression of >2,000 genes altered under each stress condition tested. Importantly, there were significant overlaps in the differentially expressed genes among the different stresses, highlighting the presence of a shared general mechanism. These genes are involved in common stress responses (e.g., translation and ribosome, tRNA, and ATP metabolic process, glycolysis, gluconeogenesis, protein geranylgeranylation, proteasome assembly, Sec61 translocon, isoprenoid biosynthesis, food vacuole, apoptosis, and mitochondrion targeting) and stress-specific response (e.g., P-bodies, protein folding, and mitochondrion targeting upon starvation, HS, and ART treatment, respectively). Especially, seven K13-interacting proteins (KICs) involved in ART resistance by participating in hemoglobin uptake [42] were down-regulated under all three stress conditions. Furthermore, KIC4 was dysregulated in PfGCN5 KD parasites during stress, indicating that the K13 regulatory pathway is a common stress response. Similarly, a large-scale forward-genetic screen in *P. falciparum* revealed apicoplast-targeted proteins including isoprenoid biosynthesis and its downstream protein modifications (geranylgeranylation and farnesylation) as a common pathway mediating tolerance to febrile temperature, a low dose of ART, and oxidative stress [4, 11]. Likewise, ART-resistant parasites showed elevated stress responses [4, 21–25]. In addition, this study also identified stress-specific responses, such as upregulation of *Pfhsp70-1* and *Pfhsp90* after HS, ATP synthesis under starvation, and ER stress response after ART treatment. These data highlight that the malaria parasite has evolved an integrated mechanism responding to different stress conditions. Intriguingly, PfGCN5 transcription was not altered upon stress treatment at the early ring stage, but the protein levels were significantly increased, indicating PfGCN5 is regulated by a post-transcriptional regulation mechanism during stress response.

In addition to the parasite-specific functions of PfGCN5 in regulating invasion and virulence in *P. falciparum*, we confirm an evolutionarily conserved role of PfGCN5 in regulating stress responses. By comparing gene expression between PfGCN5-normal and PfGCN5-deficient parasites exposed to stresses, we identified a subset of 300-400 genes whose expression in response to stresses depended on PfGCN5. These PfGCN5-dependent, stress-response genes are shared in important pathways (translation, energy metabolism, Pyrimidine and amino acid metabolic process, Sec61 translocon, Maurer’s cleft, and food vacuole) as well as in stress-condition-specific pathways (DHA: protein geranylgeranylation, regulation of protein phosphorylation/kinase, proteasome assembly, protein targeting mitochondrion, and heterochromatin; the low glucose: translation initiation, pentose-phosphate shunt, PTEX complex, response to drug and heat, and RNA process; HS: translational elongation, protein folding and response to the xenobiotic stimulus), indicating that PfGCN5 is a key activator of general and specific stress responses. Interestingly, almost all AP2 TFs were altered in the same pattern in the PfGCN5 KD under three stress conditions except that four AP2 TFs were differentially changed under HS. PfGCN5 is present in a large coactivator protein complex(es) to regulate global gene expression in *P. falciparum* [36]. Activation of specific pathways by PfGCN5 may be conferred by specific transcription factors such as the AP2-domain proteins. In the PfGCN5 complex, PfAP2-LT is present as a consistent member, and other AP2 TFs (e.g., AP2-I and PF3D7_1239200) have also been found in PfGCN5 pulldowns [36, 49]. Of particular relevance, PfAP2-HS was found to play a key role in the HS response through the activation of *Pfhsp70-1* and *Pfhsp90* [5]. While we hypothesize that the PfGCN5 complex may be dynamically recruited to activate genes in the stress response pathways by PfAP2-HS, the exact mechanism remains to be tested.

The observation of delayed recovery and lower RSA rate in response to DHA exposure in parasites with reduced levels of the PfGCN5 protein is in agreement with the general increase in stress tolerance that is usually seen in the early stages of resistance, which is a characteristic of artemisinin resistance [50]. Re-sensitization of the ART-resistant parasites by chemically inhibiting PfGCN5 may provide a way to deal with the emerging problem of ART resistance in endemic areas and underline PfGCN5 as a potential target for therapeutic development.

## MATERIAL AND METHODS

### Parasite culture

The *P. falciparum* strain 3D7 and its genetically modified clones were cultured at 37°C in a gas mixture of 5% CO_2_, 3% O_2,_ and 92% N_2_ with type O^+^ RBCs at 5% hematocrit in RPMI 1640 medium supplemented with 25 mM NaHCO_3_, 25 mM HEPES, 50 mg/L hypoxanthine, 2 g/L glucose, 0.5% Albumax II and 40 mg/ml gentamicin sulfate [51]. Synchronization of asexual stages was performed by sorbitol treatment at the rings stage followed by incubation of synchronized schizonts with fresh RBCs for 3 h to obtain highly synchronized ring-stage parasites [52].

### Genetic manipulation of *PfGCN5*

To generate a PfGCN5 KD parasite line by TetR-DOZI system, a *PfGCN5* fragment [nucleotides (nt) 3778-4758] was amplified using primers F1 (GA<u>GCGCGC</u>TGTTACCTCAACTGAGC, *BssH*II underlined) and R1 (GA<u>GGTTACC</u>TGCTGTATCAGTTATAGCTTC, *BstE*II underlined) from *P. falciparum* genomic DNA and cloned into pMG75 ATPase4 plasmid [37, 38] to replace the ATPase4 fragment and generate pMG75-PfGCN5. To fuse the C-terminal of PfGCN5 with GFP, we amplified GFP using primers F2 (CA<u>GGTTACC</u>ATGAGTAAAGGAGAAGAACTTTTC, *BstE*II underlined) and R2 (CT<u>GACGTCT</u>TATTTGTATAGTTCATCCATGCC, *Aat*II underlined) and cloned it into pMG75-PfGCN5 at the *BstE*II and *Aat*II sites.

Parasite transfection was done using the RBC loading method [53]. Briefly, 100 µg of plasmid was introduced into fresh RBCs by electroporation. Purified schizonts were used to infect the RBCs pre-loaded with the plasmid, and selection was done with blasticidin (BSD) at 2.5 μg/mL for approximately 4 weeks with weekly replenishment of fresh RBCs until resistant parasites appeared. Resistant parasites were subjected to three cycles of drug on-off selection and single clones of parasites with stable integration of the constructs were obtained by limiting dilution [52]. aTc at 0.5 μM was constantly added to the culture to maintain adequate expression of *PfGCN5*. GFP-positive parasites were sorted and cloned by fluorescence-activated cell sorting. Correct integrations of plasmids into the parasite genome were screened by Southern blot with the digoxigenin (DIG)-labeled probes using an established protocol [36, 54]. The probe was generated by using the F1 and R1 primers.

### PfGCN5 KD and growth phenotype analysis

Flow cytometry was used to measure the GFP level in the TetR-PfGCN5::GFP parasites. The growth of the TetR-PfGCN5::GFP parasite line was measured in triplicate. Cultures were tightly synchronized, as described above. The parasitemia of the culture was monitored daily by microscopy of Giemsa-stained blood smears. Growth rates after *PfGCN5* KD were analyzed by starting cultures at 0.1% rings with the vehicle control (ethanol, -aTc) or aTc (+aTc) for seven days. Growth rates after HS and low glucose treatment (6 h) were measured by starting cultures at 0.5% rings with the vehicle control (ethanol, -aTc) or aTc for five days. Parasite recovery assay was performed as described previously by treating 2% early ring-stage parasites with 1 μM of DHA for 12 h [55]. aTc was added back to the PfGCN5 KD (-aTc) culture after removing stress conditions.

### Western blot and live imaging

To assess the PfGCN5 protein expression during the IDC in normal or stress conditions, synchronized parasite cultures were lysed with 0.06% saponin, and the parasite pellet was washed thrice with PBS. Proteins were extracted by incubating parasite pellets with 2% SDS for 30 min at room temperature. Pellets were centrifuged at 10,000g for 5 min to collect supernatants. Proteins from rings, trophozoites, and schizonts were equally loaded and resolved in 4-20%SDS-PAGE. Western blot analysis was performed using TetR-PfGCN5::GFP, PfGCN5::GFP, and PfGCN5::PTP parasite lines expressing GFP or PTP-tagged PfGCN5 detected by rabbit anti-GFP (1:2000, Novus Biologicals) or rabbit anti-protein C (1:3000, GenScript) as primary antibodies, respectively [36]. HRP-conjugated goat anti-rabbit IgG (1:5000, Millipore) was used as the secondary antibody. Anti-PfH3 antibody (1:5000, Sigma) was used as the loading control. The results were visualized with the ECL detection system (Clarity Max™, Bio-Rad) and the grey values of the bands detected by Western blot were quantified using the ImageJ software.

To assess the GFP expression in live TetR-PfGCN5::GFP parasites, images were captured using a Zeiss Axiovert Microscope equipped with 100× objective after the nuclei were stained by Hoechst 33342 (20 mM, Thermofisher). Acquired images were processed using Zeiss Zen software and Adobe Photoshop 2012.

### Transcriptome analysis

To compare the parasites’ transcriptomes upon stress conditions, we performed RNA-seq analysis using the ring-stage TetR-GCN5::GFP parasites with or without aTc. The experiment was done in 2-3 replicates. Total RNA was extracted using the ZYMO RNA purification kit and RNA integrity was confirmed by the TapeStation system (Agilent). Total RNA was used to generate the sequencing libraries using the KAPA Stranded mRNA Seq kit for the Illumina sequencing platform according to the manufacturer’s protocol (KAPA biosystems). Libraries were sequenced on an Illumina NextSeq 550 using 150 nt paired-end sequencing. Reads from Illumina sequencing were mapped to the *P. falciparum* genome sequence (Genedb v3.1) using HISAT2 [56]. The expression levels and the differential expression were calculated by FeatureCounts and DESeq2 [57, 58] with the criteria of ≥ 1.5-fold alteration and *P*-adjustment <0.1. RNA-seq data were submitted to the NCBI GEO repository (accession number GSE221211) with a token (cxmngauyrvcfjqj) for reviewers’ access.

### GO enrichment analysis and clustering

The GO enrichment was performed on PlasmDB (https://plasmodb.org/plasmo/). The fold changes of gene expressions were further normalized by Z-score, and K-means were performed to identify differential gene expression patterns among the transcriptomes with or without KD of PfGCN5 under different stress conditions.

### ChIP-qPCR

ChIP-qPCR was performed as described with some modifications [36]. Synchronized TetR-PfGCN5::GFP parasite lines at the early ring stage (6–12 hpi, ∼5 × 10^9^ iRBCs) were harvested and crosslinked with 1% paraformaldehyde and then neutralized by glycine (0.125 M). The fixed iRBCs were lysed with saponin (0.06% final concentration) and parasites were treated with a lysis buffer (10 mM KCl, 0.1 mM EDTA, 0.1 mM EGTA, 1 mM DTT, 10 mM Hepes, pH 7.9, 1 × protease inhibitor) and then were gently homogenized using a douncer to free the nuclei. Pelleted nuclei were sonicated in a shearing buffer (0.1% SDS, 5 mM EDTA, 50 mM Tris-HCl, pH 8.1, 1× protease inhibitor). using a rod bioruptor (Microson ultrasonic cell disruptor, Misonix, Inc. USA) at high power for 20 cycles of 30 sec ON/30 sec OFF, resulting in sheared chromatin of approximately 100–1000 bps. Fifty μl of input samples were set aside, and the remaining chromatin was diluted in an incubation buffer (0.01% SDS, 1.5% Triton X-100, 0.5 mM EDTA, 200 mM NaCl, 5 mM Tris-HCl, pH 8.1). The chromatin (75 μl/400 ng) was incubated with rabbit anti-GFP antibodies (NB100-1770SS, Novus biologicals) overnight at 4°C while rotating followed by the addition of 20 μl of agarose beads. Beads were then washed with the following: buffer 1 (0.1% SDS, 1% Triton X-100, 150 mM NaCl, 2 mM EDTA, 20 mM Tris HCl, pH 8.1); buffer 2 (0.1% SDS, 1% Triton X-100, 500 mM NaCl, 2 mM EDTA, 20 mM Tris HCl, pH 8.1), buffer 3 (250 mM LiCl, 1% NP-40, 1% Na-deoxycholate, 1 mM EDTA, 10 mM Tris HCl, pH 8.1) and finally twice with buffer 4 (10 mM EDTA, 10 mM Tris HCl, pH 8). The immunoprecipitated (IPed) chromatin was eluted with the elution buffer (1% SDS, 0.1 M NaHCO_3_) at room temperature. The eluted chromatin and input samples were reverse crosslinked and purified by the phenol:chloroform method. For qPCR, 10 ng per well in triplicate was used with the FastStart Universal SYBR Green Master [Rox] (Sigma-Aldrich, USA). Primer pairs targeting 5’ UTRs of the selected genes were designed to amplify fragments less than 300 bp (**Table S5**). Fold enrichment relative to constitutively expressed reference gene *seryl-tRNA synthetase* was calculated using the 2^−ΔΔCt^ method. The fold changes of binding enrichment were calculated using a formula: 2− [(IP Ct-target −IP Ct-stRNA) − (Input Ct-target–Input Ct-stRNA)] for each primer set targeting specific promoter regions.

### *In vitro* drug assay, recovery assays after DHA treatment, and RSA

The standard SYBR Green I-based fluorescence assay [59, 60] was used to assess parasite susceptibilities to MB-3. Synchronized cultures at the ring stage were diluted with fresh complete medium to 1% hematocrit and 0.5% parasitemia. *In vitro* drug assays were performed in 96-well microtiter plates with serially diluted drug concentrations. Three technical and biological replications were performed. The recovery assay was performed based on established methods with some modifications [25, 55, 61–63]. Briefly, the highly synchronized early ring stage (0-6h) parasites at 2% parasitemia were treated with 1 μM DHA for 12 h. The parasite growth was examined daily using Giemsa staining. The amount of time needed for each parasite culture to reach 5% parasitemia was recorded. RSA was performed as previously described [41, 60, 64–66]. Briefly, schizonts were purified from tightly synchronized cultures using a Percoll gradient and allowed to rupture and invade fresh RBCs for 3 h. The cultures were treated with sorbitol to select early rings and eliminate the remaining schizonts. Ring-stage parasites of 0-3 hpi at 1% parasitemia and 1% hematocrit were exposed to 700 nM DHA for 6 h, followed by a single wash. MB-3 was added and washed simultaneously as DHA, and aTc was added back to the PfGCN5 KD (-aTc) culture after drug treatment. After culturing for 66 h, ∼10,000 RBCs were observed on thin blood smears to count viable parasites.

### Statistical analysis

For all experiments, three or more independent biological replicates were performed. The results were presented as mean ± SD. Results were regarded as significant if *P* < 0.05 as established by T-test, and the respective analysis was shown in the figure legends.

## Acknowledgments

This study was supported by the startup fund from Morsani College of Medicine, University of South Florida, and grant R21AI149202 from the National Institute of Allergy and Infectious Diseases (NIAID), NIH, USA to JM. We are grateful to Jacobus *Pharmaceuticals* for providing the drug WR99210. We thank the University of South Florida Genomics Program (Sequencing Core and Computational Core/Omics Hub) for supporting NGS sequence and data analysis.

## Author Contributions

J.M. conceived and designed the study. AR.S, C.W., and AB.L. performed research, acquired and analyzed data. C.W. analyzed the RNA-seq data. M.K. and A.C. assisted research. X.L. conducted parasite culture and phenotypic growth analysis. J.M. wrote the original draft.

---

**Figure S1. PfGCN5 knockdown using the TetR-DOZI system**. **A.** Schematic diagram shows the insert of 10 x aptamer (A*) in 3’ UTR of *PfGCN5* and integration of the TetR-DOZI expression cassette by single-crossover homologous recombination. KAT, lysine acetyltransferase domain; ADA2, ADA2-binding domain; BrD, bromodomain; H, HindIII restriction site. The recombined PfGCN5 locus includes GFP tagging at the C-terminus of PfGCN5. **B.** Southern blot indicates three positive clones from transfected parasites. M, molecular markers in Kb. **C.** Western blot (left panel) showed that PfGCN5-GFP level and its processed fragments in TetR-PfGCN5::GFP compared to PfGCN5::GFP. The histone H3 was used as a loading control. The relative intensity of PfGCN5 was measured by densitometry (right panel). **D.** Recovery of PfGCN5-GFP expression after adding aTc back to the KD parasite culture for 2-10h by detecting the GFP level in the parasites via flow cytometry. The KD parasites were cultured without aTc (-aTc) for five IDCs before adding aTc. The median GFP levels of 5000 parasites were used for each replicate. The percentage of GFP level compared to parasite cultured without withdrawal of aTc (+aTc). **E**. Western blots detected the recovery of PfGCN5-GFP expression after adding aTc back to the KD parasite culture for 2-10h compared to the same parasites with (+) or without (−) aTc (left panel). The histone H3 was used as a loading control. The relative intensity of PfGCN5-GFP was measured by densitometry (right panel). The KD parasites were cultured without aTc (-aTc) for five IDCs before adding aTc. **F**. The growth rates of parasites without KD of PfGCN5 (+aTc) and parasites with re-stored PfGCN5 expression by adding aTc back to the −aTc parasites (+aTc post KD).

**Figure S2. Induction of PfGCN5 expression upon stress treatments and recovery assay. A.** The left panel shows the changes of PfGCN5-GFP expression after +aTc TetR-PfGCN5:GFP parasites were treated with heat shock (HS, 41 °C), low-glucose (LG, 0.5 g/L), and DHA (30mM) for 6 h at the late stage (trophozoite, 24-30 hpi) by Western blot. The histone H3 was used as a loading control. The right panel indicates the relative intensity of full-length PfGCN5-GFP bands among the parasites with or without stress treatments by densitometry. **B**. Recovery assays of 3D7 parasites with (+aTc) or without (−aTc) aTc after 1 μM DHA treatment.

**Figure S3. Drastic transcriptional changes upon stress conditions**. **A.** Principal component analysis (PCA) for the RNA-seq samples shows high consistency between biological replicates. Dots in the same color represent different biological replicates in the same treatment. +DHA, +HS and +Glu denote DHA (30 nM), HS (41°C), and low glucose (0.5 g/L) treatment for 6h, respectively. +aTc and –aTc indicate TetR-PfGCN5::GFP parasites were cultured with aTc and without aTc (knockdown). **B** and **C**. Heatmaps display the GO enrichment analyses of up-(**B**) and down-(**C**) regulated genes upon HS and low glucose (L-Glu), and DHA treatments based on the cellular component (CC) showing the common and stress-specific stress responses. PDC: pyruvate dehydrogenase complex, IMC: inner membrane pellicle complex, TCA: mitochondrial tricarboxylic acid cycle enzyme complex, PKA complex: cAMP-dependent protein kinase complex, Mpp10 complex (ribosome biogenesis). **D**. Heatmap shows the fold change of AP2 TFs expression under different stress conditions compared to the gene expression under no stress conditions. The arrows indicate the four AP2 TFs were up-regulated under DHA and low-glucose conditions but downregulated under HS.

**Figure S4. GO enrichment analysis and overlaps among the genes in cluster I-V. A.** GO enrichment analyses of genes in cluster III-V. The GO terms shared among the three stress conditions are shown in bold and black color, while stress-specific enrichments are listed by words in red, blue, and green for HS, low glucose, and DHA treatments, respectively. **B.** Overlapping pie charts show the number of genes in each cluster and the levels of overlaps among genes in clusters I-V upon DHA, HS, and low-glucose treatments, respectively.

**Table S1.** The transcriptomic changes in +aTc TetR-PfGCN5:GFP parasites at the ring stage under stress conditions.

**Table S2.** The transcriptomic changes in TetR-PfGCN5:GFP parasites at the ring stage after KD of PfGCN5.

**Table S3**. The transcriptomic changes in -aTc TetR-PfGCN5:GFP parasites at the ring stage under stress conditions.

**Table S4.** Expression patterns by comparing -aTc and +aTc TetR-PfGCN5:GFP parasites under three stress conditions.

**Table S5**. The primers used in this study.

