## Supplementary material for "*Plasmodium falciparum* GCN5 plays a key role in regulating artemisinin resistance–related stress responses": Figure S1-S4

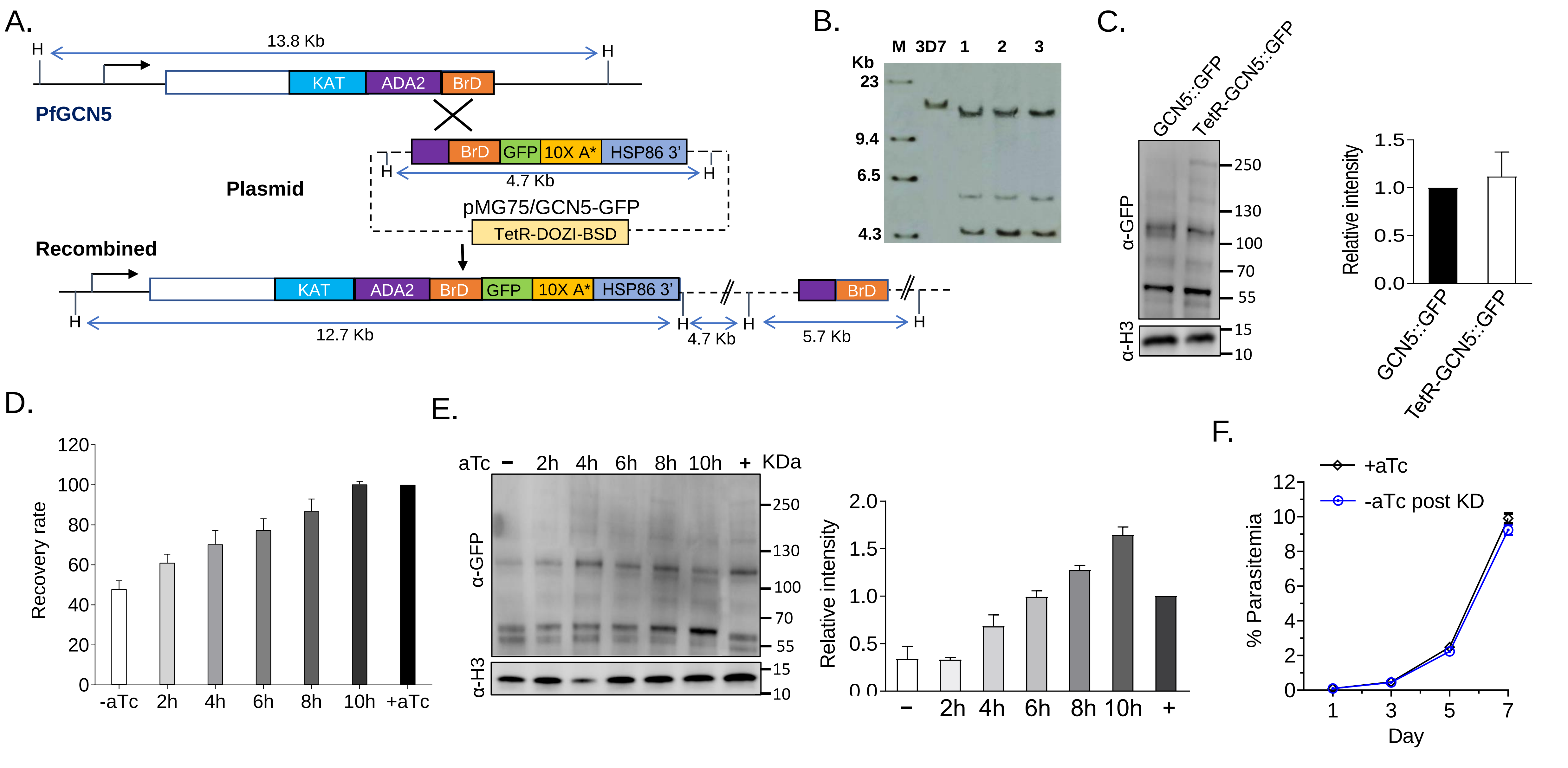


**Figure S1. PfGCN5 knockdown using the TetR-DOZI system. A.** Schematic diagram shows the insert of 10 x aptamer (A*) in 3' UTR of *PfGCN5* and integration of the TetR-DOZI expression cassette by single-crossover homologous recombination. KAT, lysine acetyltransferase domain; ADA2, ADA2-binding domain; BrD, bromodomain; H, HindIII restriction site. The recombined PfGCN5 locus includes GFP tagging at the C-terminus of PfGCN5. **B.** Southern blot indicates three positive clones from transfected parasites. M, molecular markers in Kb. **C.** Western blot (left panel) showed that PfGCN5-GFP level and its processed fragments in TetR-PfGCN5::GFP compared to PfGCN5::GFP. The histone H3 was used as loading control. The relative intensity of PfGCN5 was measured by densitometry (right panel). **D.** Recovery of PfGCN5-GFP expression after adding aTc back to the KD parasite culture for 2-10h by detecting the GFP level in the parasites via flow-cytometry. The KD parasites were cultured without aTc (-aTc) for 5 IDCs before adding aTc. The median GFP levels of 5000 parasites were used for each replicate. The percentage of GFP level compared to parasite cultured without withdrawal of aTc (+aTc). **E.** Western blots detected the recovery of PfGCN5-GFP expression after adding aTc back to the KD parasite culture for 2-10h compared to the same parasites with (+) or without (−) aTc (left panel). The KD parasites were cultured without aTc (-aTc) for 5 IDCs before adding aTc. The histone H3 was used as a loading control. The relative intensity of PfGCN5-GFP was measured by densitometry (right panel). **F**. The growth rates of parasites without KD of PfGCN5 (+aTc) and parasites with re-stored PfGCN5 expression by adding aTc back to the -aTc parasites (+aTc post KD).

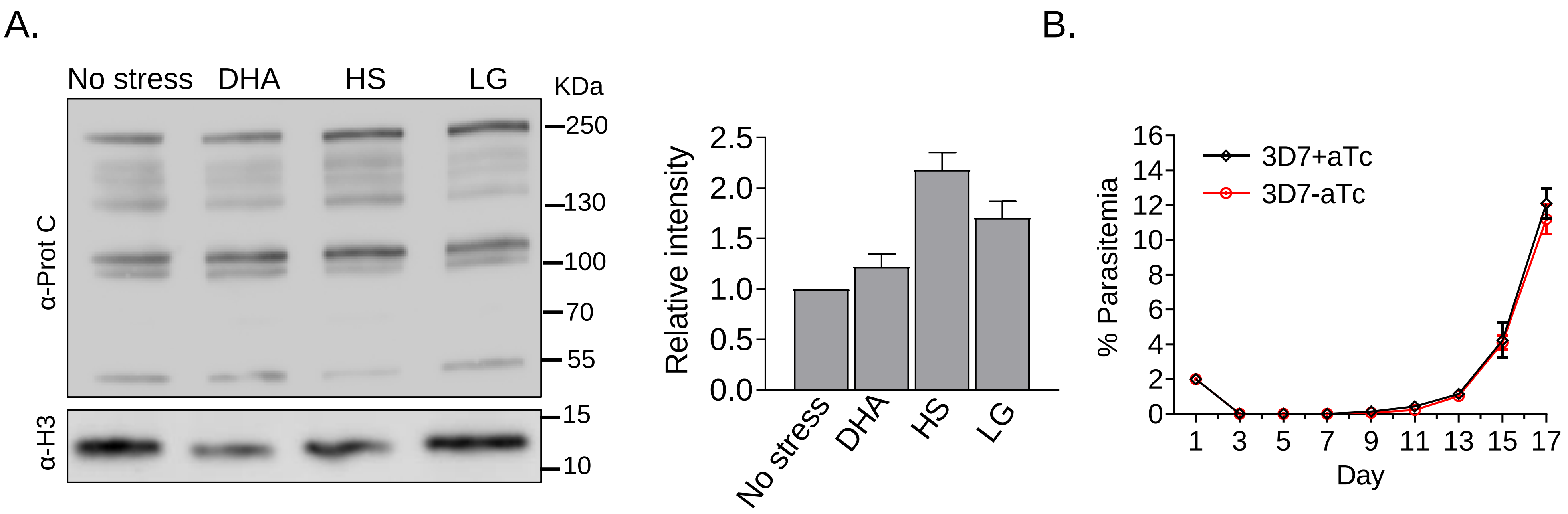


**Figure S2. Induction of PfGCN5 expression upon stress treatments and recovery assay. A.** The left panel shows the changes of PfGCN5-GFP expression after +aTc TetR-PfGCN5:GFP parasites were treated with heat shock (HS, 41 $^{o}$C), low-glucose (LG, 0.5 g/L), and DHA (30mM) for 6 h at the late stage (trophozoite, 24-30 hpi) by Western blot. The histone H3 was used as a loading control. The right panel indicates the relative intensity of full-length PfGCN5-GFP bands among the parasites with or without stress treatments by densitometry. **B**. Recovery assays of 3D7 parasites with (+aTc) or without (−aTc) aTc after 1 μM DHA treatment.

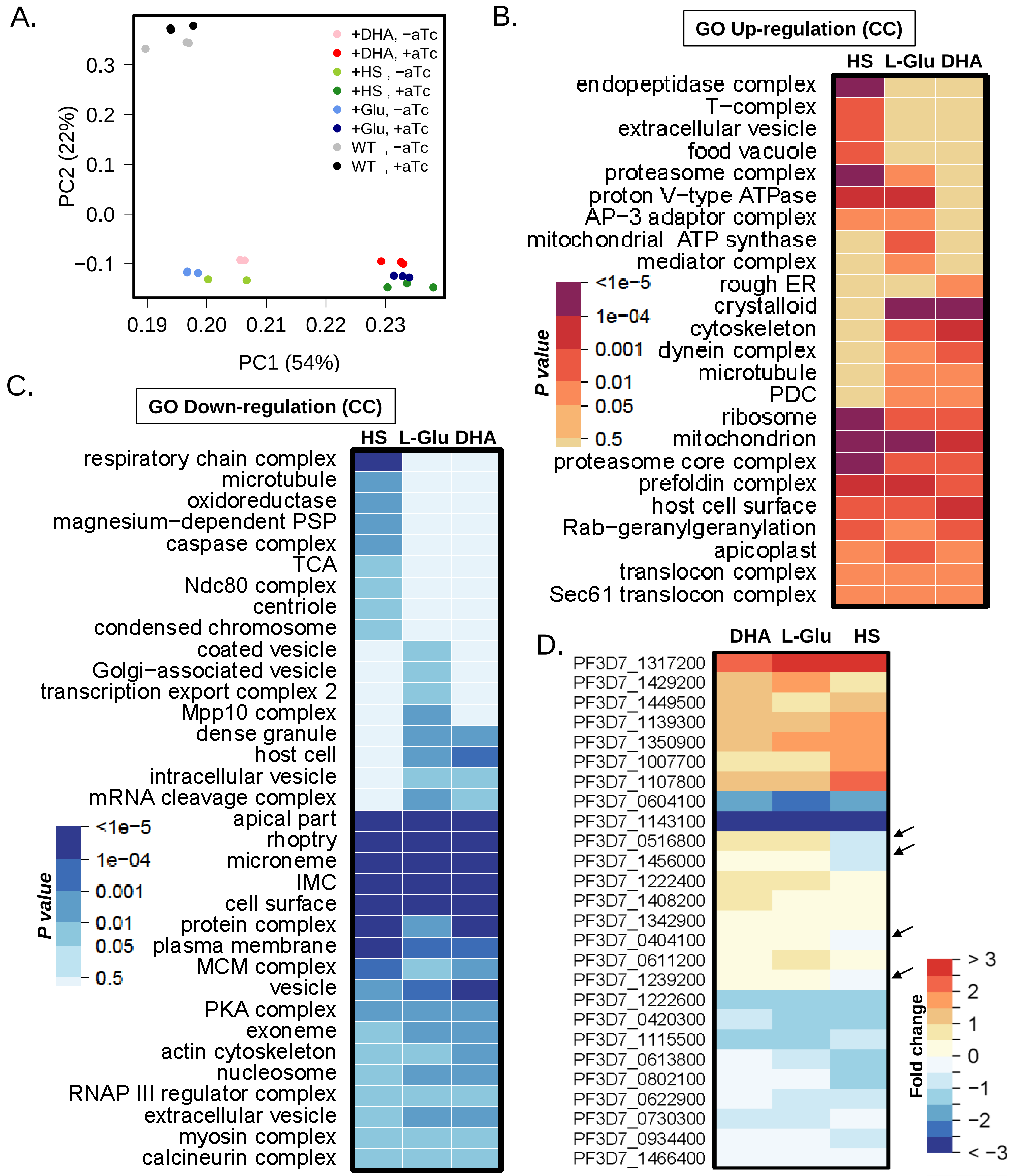


**Figure S3. Drastic transcriptional changes upon stress conditions**. **A.** Principal component analysis (PCA) for the RNA-seq samples shows high consistency between biological replicates. Dots in the same color represent different biological replicates in the same treatment. +DHA, +HS and +Glu denote DHA (30 nM), HS (41$^{o}$C), and low glucose (0.5 g/L) treatment for 6h, respectively. +aTc and –aTc indicate TetR-PfGCN5::GFP parasites were cultured with aTc and without aTc (knockdown). **B** and **C**. Heatmaps display the GO enrichment analyses of up- (**B**) and down-(**C**) regulated genes upon HS and low-glucose (L-Glu), and DHA treatments based on the cellular component (CC) showing the common and stress-specific stress responses. PDC: pyruvate dehydrogenase complex, IMC: inner membrane pellicle complex, TCA: mitochondrial tricarboxylic acid cycle enzyme complex, PKA complex: cAMP-dependent protein kinase complex, Mpp10 complex (ribosome biogenesis). **D**. Heatmap shows the fold change of AP2 TFs expression under different stress conditions compared to the gene expression under no stress conditions. The arrows indicate the four AP2 TFs were up-regulated under DHA and low-glucose conditions but downregulated under HS.

A.

Heat shock, Low glucose, DHA treatment

| Cluster | |
|---|---|
| III | **DNA replication and repair, cell cycle, isoprenoid biosynthesis, SERA**<br>Cell adhesion, cytoskeleton organization,<br>Mitochondrion electron transport |
| IV | **Proteasome, protein folding, mitochondrion, apicoplast, stress granule assembly, cytoskeleton organization, transport, fatty acid synthesis, gametocyte proteomics, ERAD**<br>Response to heat and oxidative stress, ERAD, tRNA, amino acid and vitamin B6 metabolism, protein geranylgeranylation, microtube-based movement, cation homeostasis and pH reduction<br>Response to ER stress and abiotic stimulus, cell redox homeostasis, cell cycle |
| V | **Entry into and exit from host, actin filament-based movement, endocytosis, protein modifications, food vacuole**<br>Response to drug, lipid metabolic process<br>Response to starvation, mRNA process, vesicle organization<br>Response to drug and xenobiotic stimulus |

B.

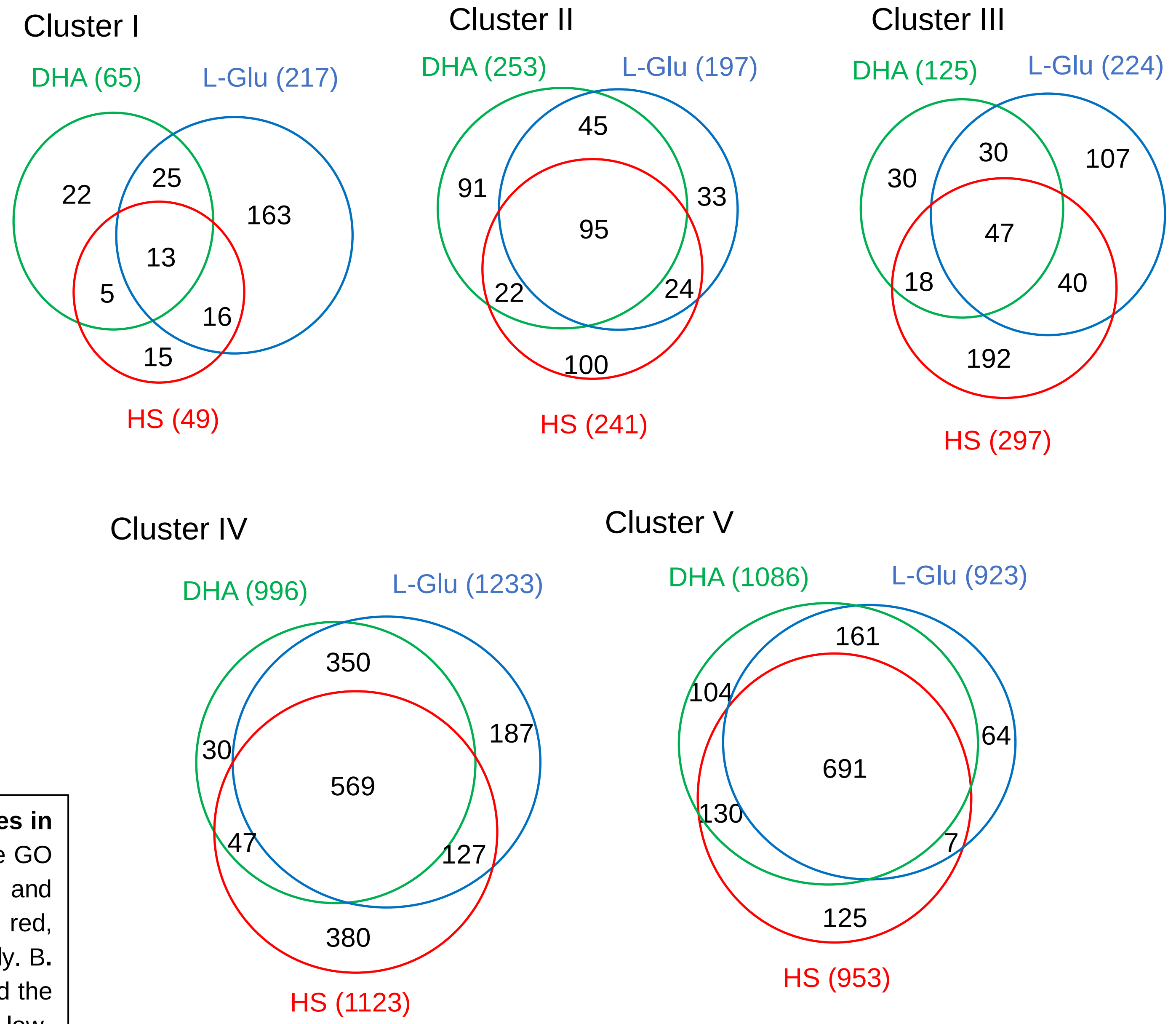


**Figure S4. GO enrichment analysis and overlaps among the genes in cluster I-V. A.** GO enrichment analyses of genes in cluster III-V. The GO terms shared among the three stress conditions are shown in bold and black color, while stress-specific enrichments are listed by words in red, blue, and green for HS, low glucose, and DHA treatments, respectively. B**.** Overlapping pie charts show the number of genes in each cluster and the levels of overlaps among genes in clusters I-V upon DHA, HS, and low-glucose treatments, respectively.
